# Implicit Learning of Temporal Behavior in Complex Dynamic Environments

**DOI:** 10.1101/2020.01.29.924472

**Authors:** Josh M. Salet, Wouter Kruijne, Hedderik van Rijn

**Affiliations:** Department of Experimental Psychology, University of Groningen

## Abstract

Humans can automatically detect and learn to exploit repeated aspects (regularities) of the environment. Timing research suggests that such learning is not only used to anticipate what will happen, but also when. However, in timing experiments, the intervals to be timed are presented in isolation from other stimuli and explicitly cued, contrasting with naturalistic environments in which intervals are embedded in a constant stream of events and individuals are hardly aware of them. It is unclear whether laboratory findings from timing research translate to a more ecologically valid, implicit environment. Here we show in a game-like experiment, specifically designed to measure naturalistic behavior, that participants implicitly use regular intervals to anticipate future events, even when these intervals are constantly interrupted by irregular yet behaviorally relevant events. This finding extends previous research by showing that individuals not only detect such regularities but can also use this knowledge to decide when to act in a complex environment. Furthermore, this finding demonstrates that this type of learning can occur independently from the ordinal sequence of motor actions, which contrasts this work with earlier motor learning studies. Taken together, our results demonstrate that regularities in the time between events are implicitly monitored and used to predict and act on what happens when, thereby showing that laboratory findings from timing research can generalize to naturalistic environments. Additionally, with the development of our game-like experiment we demonstrate an approach to test cognitive theories in less controlled, ecologically more valid environments.

---

Implicit Learning of Temporal Behavior in Complex Dynamic Environments While the brain is continuously confronted with a highly dynamic stream of information, this sensory flood contains regularities which the brain capitalizes upon to reduce processing load. For example, regularities in stimulus sequences are learned (Brady & Oliva, 2008; Fiser & Aslin, 2002; Turk-Browne, Jungé, & Scholl, 2005), can bias which objects are encoded in working memory (Umemoto, Scolari, Vogel, & Awh, 2010), and can guide attention (Baker, Olson, & Behrmann, 2004; Olson & Chun, 2001; Yu & Zhao, 2015; Zhao, Al-Aidroos, & Turk-Browne, 2013). Such ‘statistical learning’ often takes place automatically and implicitly. By automatically extracting regularities from the environment, observers form a compact representation of the world that, according to the predictive coding framework (Clark, 2013; Friston, 2005; Mehta, 2001), can be used to generate predictions about future events.

It has been suggested that observers also detect and utilize temporal regularities in the environment, in order to anticipate not only what will happen, but also when. A long tradition of timing research has shown that regularities in the interval duration between events can be learned and steer preparatory behavior: conditioned rabbits close their eyes in response to an air puff, but only do so around the point in time of expected delivery (Schneiderman & Gormezano, 1964); infants automatically track the interval duration of rhythmic auditory stimuli (Brannon, Roussel, Meck, & Woldorff, 2004); and adults optimize preparation at time points where an event is likely to occur (Coull & Nobre, 1998; Niemi & Näätänen, 1981; Steinborn, Rolke, Bratzke, & Ulrich, 2009; Woodrow, 1914). Los, Kruijne, and Meeter (2014) argued that such adaptation to regular intervals is driven by an implicit, automatic memory process. Recently, we have implemented this memory proposal in a computational framework and showed that adaptation effects to the distribution of intervals used in the experiment can be accounted for by automatic Hebbian learning between a representation of time and a motor layer (Salet, Kruijne, Van Rijn, Los, and Meeter, under review). This further support the view that adaptation to regular intervals stem from implicit mechanisms, much like statistical learning.

However, in typical timing studies the intervals to be timed are isolated in single trials in which their onset and offset are explicitly cued, and therefore participants are well aware of the importance of temporal information for performance. This transparency contrasts starkly with naturalistic environments in which individuals are rarely aware of intervals in constant streams of both regular and irregular events. Theoretically, this contrast is important for understanding temporal behavior because identifying and utilizing temporal regularities in the world might be much more difficult. An important question is therefore whether laboratory findings from timing studies translate to a more ecologically valid, implicit environment. Do individuals still learn and adapt to regular intervals when they are unnoticed and interrupted by other irregular events?

While implicit learning of temporal regularities has received considerable attention in the statistical learning literature, it remains unclear whether implicit knowledge about regular intervals can be used to optimize preparatory behavior. One line of work (e.g., Brady & Oliva, 2008; Kidd, Piantadosi, & Aslin, 2012; Turk-Browne et al., 2005), studied adaptation to the regularity formed by their order (e.g., stimulus ‘B’ always follows stimulus ‘A’) and not by interval duration (e.g., ‘B’ reappears after a predictable interval, yet can be followed by either ‘A’ or ‘C’). Other studies have shown that regular intervals can be implicitly learned: infants (Lewkowicz, 2003) and adults (Damsma and van Rijn, 2017, Olson and Chun, 2001, Experiment 1a and b) can implicitly detect rhythmic interval patterns; temporal regularities influence the learning of language (Hay & Saffran, 2012) and artificial pitch grammar (Selchenkova, Jones, & Tillmann, 2014); temporal structure in a rapid serial visual presentation task can bias the probability of shifting the focus of attention (Sali, Anderson, & Yantis, 2015); and temporal information can be integrated in the learning of rhythmic perceptual-motor action sequences (Gobel, Sanchez, & Reber, 2011; O’Reilly, McCarthy, Capizzi, & Nobre, 2008; Schultz, Stevens, Keller, & Tillmann, 2013; Shin & Ivry, 2002). However, it is unclear whether implicit knowledge about regular intervals is used to anticipate when to act. Here, we aim to demonstrate that, similar to reports from timing studies mentioned in the previous paragraphs, individuals both detect and act upon regular temporal information to optimize preparation at specific time points. Additionally, we aim to test implicit learning in a complex dynamic environment where, in contrast to typical studies on implicit learning, regular events are continuously interrupted by irregular yet behaviorally relevant events.

To this end, we developed a novel paradigm inspired by the arcade game ‘Whac-A-Mole’. Participants scored points by ‘hitting’ sudden-onset targets (the ‘moles’) using the mouse. At two out of the three target positions targets appeared at random times, but at one position targets consistently appeared every three seconds. We analyzed performance expressed by response times, hit rates, and participants’ cursor movements in epochs around target onsets. Hypothesizing that participants adapt to the temporal regularity, we expect better performance for regular than for irregular targets. Additionally, we split the experiment into an explicit and an implicit phase, where participants respectively were and were not informed about the regularity. This allowed us to further investigate whether temporal regularities only drive behavior when they are explicitly signalled beforehand, or whether they can also be implicitly detected and acted upon without prior information.

## Experiment 1

### Method

#### Participants

Forty-eight first-year psychology students of the University of Groningen were recruited and received course credit for their participation. Sample size was motivated on the basis of previous studies that have used reaction time (RT) as a measure of implicit learning (Zhao et al., 2013): a power analysis suggested a sample size of 39 participants to detect a reliably difference between regular and irregular targets (Cohens’s *d* = 0.46, power = 80 %, *α* = 0.05). We tested 48 participants to fully counterbalance the location of the regular target across experimental conditions. The experimental procedure was approved by the Ethical committee of the Faculty of Behavioural and Social Sciences (18018-P), and participants provided written informed consent before participation.

#### Stimuli and task

The experiment was implemented using OpenSesame (Mathôt, Schreij, & Theeuwes, 2012) version 3.2.8 with the PsychoPy back-end (Peirce et al., 2019). Participants were seated in a dimly lit room at approximately 60 cm viewing distance from a 22” CRT monitor (100 Hz, Iiyama MA203DT) and were instructed to score as many points as possible by moving the cursor towards sudden-onset targets. Three light blue, unfilled circles (r = 1.5 cm) placed at the three corners of an equilateral triangle (sides: 24 cm) marked the positions of possible targets (Figure 1). A white fixation dot (r = 0.24 cm) was presented centrally. Target onsets were presented by filling the circles in with light blue for 0.5 s. In each block, targets were presented 20 times at each of the three locations. One of the locations was the ‘regular target location’ at which a target appeared every three seconds. At the other two irregular locations, targets appeared at pseudorandomly generated intervals, ensuring that the interval between two subsequent presentations in the same location was minimally 0.75 s (0.25 s between offset and new onset) and that there was always at least one irregular target between two regular target presentations.^1^ Importantly, the order of target presentation was random.

**Figure 1.**
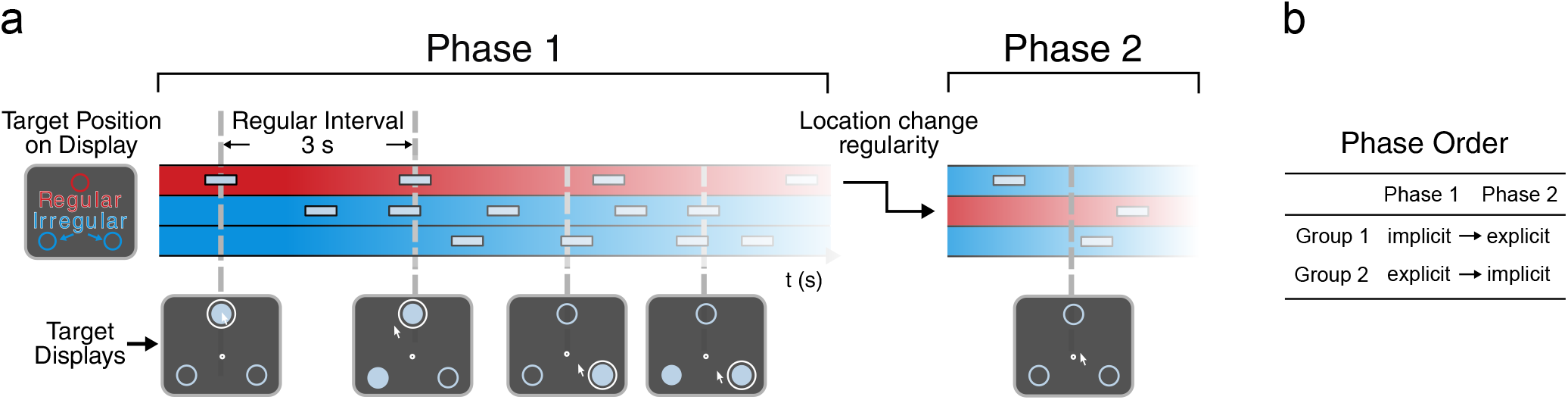
Segments of exemplary stimulus sequences in Phase 1 and Phase 2. a) The three colored rows in the chart represent the three possible target locations: at one location (red row) the regular target appears every three seconds and in the other two locations (blue rows) targets appear at pseudorandomly determined moments. Example ‘Target Displays’ are depicted at five different time points (bottom figure). Note the location change of the regularity between the two phases. b) In each of the two phases, the regularity was either implicit or explicit. The phase order was counterbalanced across participants: one participant group (‘implicit → explicit’) started with the implicit phase and the other group (‘explicit → implicit’) with the explicit phase. Colors and sizes of example displays are illustrative and do not match the experiment exactly. An example video of the experiment is available on OSF (https://tinyurl.com/w3ocv8x).

Whenever participants moved the cursor within 3 cm of one of the target locations a white circle appeared around it indicating a hit (Figure 1). Participants scored points if they hit a target within 0.5 s after its onset. Each hit was rewarded with five points and accompanied by a positively valued sound of a dropping coin. If participants were too late (0.25 s after target offset) or ‘hit’ a location that did not contain a target, three points were deducted as a penalty.

#### Procedure

The experiment consisted of two phases of eight blocks, with 60 target presentations in total. A block lasted approximately one minute, and was followed by feedback on the total score, the number of hits and misses per block, and the participants’ high score. After the first phase, the temporal regularity was assigned to a new location (Figure 1). In each of the two phases, the regularity was either implicit or explicit, and the phase order was counterbalanced across participants. One group (phase order: ‘implicit → explicit’) started with the implicit phase and was initially not informed about any regularity. Before the onset of the explicit phase, participants were fully informed about the location and timing of the regular target. They were encouraged to make use of this information, but to not ignore the targets at the other two locations. The other participant group (phase order: ‘explicit → implicit’) was informed about the regular target at the start of the experiment. Before the second (implicit) phase, they were informed that the former regularity would no longer hold. The location of the temporal regularity always switched between the first and second phase and was counterbalanced between participants. Both phases started with two practice blocks. In the explicit phase, regular targets were colored red during practice, and at the start of each block the regular location was colored red, up to the moment of first target appearance.

#### Questionnaire

After the implicit phase, participants answered three questions to assess their awareness of the temporal regularity: (1) an open question in which participants were asked whether they had observed anything “noteworthy” during the experiment, (2) after being informed of the existence of the temporal regularity, a binary question whether they had noticed this regularity, and (3) a forced choice question to indicate (or guess) the location of the temporal regularity. The complete questions are provided in Figure A1 of the Appendix.

#### Inclusion criteria

Based on the questionnaire, we excluded the data from participants that indicated to have been aware of the regularity in the implicit phase. In the open question, none of the participants reported on the regularity, but we excluded the data from two participants who indicated to have been aware of the regularity in the binary question and chose the correct location in the forced choice question (n = 46). We additionally excluded data from blocks where the hit rate of one of the three targets was less than 25 %, suggesting that a participant might not have divided its attention across all targets. For six participants, more than three out of eight blocks in either phase were discarded which led us to exclude their data entirely. Of the remaining participants (n = 40) we excluded responses faster than 0.15 s (1.2 % of cases), as well as responses to targets that appeared consecutively at the same location (9.6 % of cases), as due to the design such repetitions could only occur for irregular targets.^2^

#### Analyses

Analyses were conducted in R 3.6.3 (R Core Team, 2020). Analysis scripts are available on OSF (https://tinyurl.com/w3ocv8x). The analyses assessed whether participants adapted their behavior to the regular targets, and whether this was modulated by awareness of the regularity (explicit versus implicit). To this end, we analyzed performance expressed by RT and hit rates, and analyzed participants’ cursor movements around target onsets.

We analyzed RT by means of Linear Mixed Models (LMMs; Baayen, Davidson, & Bates, 2008; Bates, Mächler, Bolker, & Walker, 2015). We log-transformed RT to reduce the typical skew in RT distributions. Hit rates were analyzed with general (G)LMMs with a logistic link function. For both RT and hit rate models we included ‘participant’ as a random intercept term, and we used model comparisons by means of Bayes Factors (BF) estimated from BIC values (Wagenmakers, 2007) to assess whether the addition of predictors as fixed effects or random slopes was warranted.

The main interest of the study was whether regularity of targets (regular versus irregular), phase (explicit versus implicit), and their possible interaction affected performance. To investigate order effects, we assessed whether these main predictors interacted with phase order (‘explicit → implicit’ versus ‘implicit → explicit’). We additionally assessed the effect of response type on the preceding target (hit versus miss) and practice effects (time-on-task, indexed by trial number). Finally, for each experimental phase, we investigated whether the time course, indexed by experimental block (as continuous predictor), affected adaptation by testing for the interaction between experimental phase and block. Does the adaptation effect grow or shrink throughout each experimental phase? The Supplemental Materials offer a detailed description of the construction of the statistical models.

In the Results, we only report the statistics on the effects of ‘regularity’, ‘phase’, their interaction, and the time course on adaptation. The analysis of ‘phase order’ did not reveal any effect on either RT or hit rate (see Supplemental Materials). To quantify statistical evidence for or against an effect of interest, we chose to always compare the best statistical model (in terms of *BIC*) with this effect, against the best model without this predictor term. Note that for some tests, this means that the compared models can differ in terms of other predictors as well. For completeness, we also report the outcome of Likelihood Ratio Tests for these comparisons indexed by *χ*^2^ and *p*-values. But note that we relied on the *BIC*, as a prespecified criterion, for our conclusions. The Supplemental Materials offer a complete report of the statistical procedure and an overview of the models that were compared to quantify statistical evidence for *(BF*) or against (1/*BF*) an effect’s inclusion is provided on our OSF repository (https://tinyurl.com/w3ocv8x). Additionally, in the Supplemental Materials, we report traditional repeated measures analyses of variance on condition averages (ANOVA; including standardized effect sizes) to facilitate comparison with other studies.

To analyze cursor movements we selected data epochs in a time window from 0.5 s before to 1.0 s after each target onset, and calculated the distance of the cursor to the target every 0.01 s. As we did not find support for asymmetries nor carryover effects between the two phase orders, we collapsed over ‘phase order’ and subsequently assessed the effects of ‘regularity’ and ‘phase’ by means of two-way repeated measures ANOVAs at each timepoint, and identified significant differences using cluster-based permutation testing (5000 iterations, *α* = 0.05; Maris & Oostenveld, 2007), using the ‘permuco’ R package (Frossard & Renaud, 2019).

## Results

### Questionnaire

None of the participants gave any indication of awareness of the regularity on the first, open question. In the yes/no forced choice question, six out of 48 participants (12.5 %) indicated to be aware of the regularity. In the identification question, 18 participants (37.5 %) correctly chose the location of the regular target out of the three options. Two of these participants also indicated to have noticed the regularity and were omitted from further analysis.

### Reaction times

Figure 2a depicts mean RT in each block, split on ‘regularity’, ‘phase’, and ‘phase order’. Model comparisons revealed that RT to the regular target was lower than to irregular targets (Δ*BIC* = 77.40, *BF* > 1000; *χ*^2^(3) = 108.25, *p* < 0.001), lower in the explicit phase compared to the implicit phase *(ΔBIC* = 78.51, *BF* > 1000; *χ*^2^(2) = 99.08, *p* < 0.001), and that there was an interaction between ‘regularity’ and ‘phase’ *(ΔBIC* = 47.22, *BF* > 1000; *χ*^2^(1) = 57.50, *p* < 0.001), entailing that regular and irregular targets differed more in the explicit than the implicit phase. Post hoc Tukey’s HSD tests revealed that RTs to the regular target were significantly lower in both the explicit *(z* = 10.75, *p* < 0.001) and implicit *(z* = 5.18, *p* < 0.001) phase. Finally, we found no evidence that the adaptation effect grew or shrank throughout an experimental phase (Δ*BIC* = 19.75, *1/BF* > 1000; *X*^2^(4) = 5.88, *p* = 0.21).

**Figure 2.**
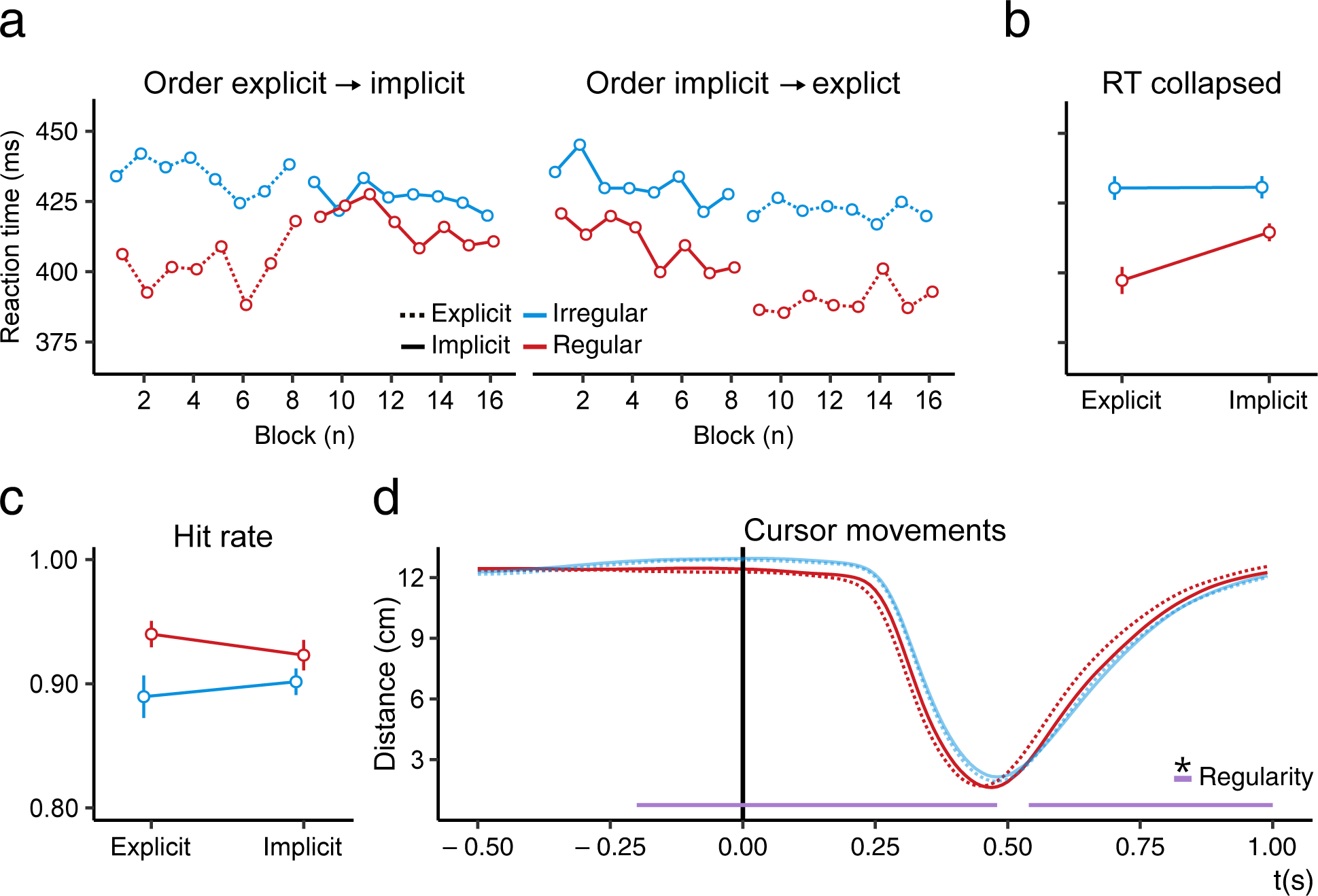
Results of Experiment 1. a) Average response times per block for regular (red) and irregular (blue) trials in the explicit (dashed lines) and implicit phase (solid lines), for phase order: ‘explicit → implicit’ and ‘implicit → explicit’. Within subject confidence intervals are omitted here as not all blocks were included for all participants (see inclusion criteria). b) Average response time collapsed over ‘phase order’ and experimental block. c) Similar to b, but for hit rates. Error bars in b and c represent 95 % within-subject confidence intervals (Cousineau, 2005; Morey, 2008). d) Average cursor distance to targets in a 0.5 to 1.0 s window around targets’ onset, for regular and irregular targets in the explicit and implicit phase. The horizontal bars at the bottom indicate significant clusters for the main effect ‘Regularity’.

### Hit rate

Figure 2c depicts the average hit rate collapsed over ‘phase order’ and experimental block. Model comparison revealed that hit rates were higher for the regular target than irregular targets *(ΔBIC* = 5.30, *BF* = 14.17; *χ*^2^(1) = 15.67, *p* < 0.001). We did not find evidence for the effect of ‘phase’ *(ΔBIC* = 10.35, 1/*BF* = 177.18; *χ*^2^(1) = 0.02, *p* = 0.90). Evidence for their interaction, indicating that the difference between the regular and irregular targets differed more in the explicit than the implicit phase, was inconclusive *(ΔBIC* = 0.02, 1/*BF* = 1.01; *χ*^2^(2) = 20.72, *p* < 0.001). Consistent with our RT analyses, model comparisons did not reveal support for an effect of time course on adaptation (Δ*BIC* = 23.75, *1/BF* > 1000; *χ*^2^(7) = 21.10, *p* = 0.003).

### Cursor movements

Figure 2c displays the average cursor movements in a window of 0.5 s before to 1.0 s after target onset, split on ‘regularity’ and ‘phase’. We found two significant clusters for the main effect ‘regularity’ (*p* < 0.001): one from 0.2 s before to 0.48 s after target onset and one from 0.54 s to the end of the epoch, both demonstrating earlier movement towards regular targets compared to irregular targets.

## Discussion

Only two out of 48 participants could accurately report on the temporal regularity in the implicit phase. In the remaining unaware participants, RT was lower, hit rate was higher, and cursor movements to the regular target were initiated earlier compared to irregular targets, as if participants anticipated the upcoming event. We found no support that such adaptation gradually developed over the course of the implicit phase, suggesting that participants detected and adapted to the regularity early in the implicit phase. Moreover, the absence of an effect of ‘phase order’ ruled out the possibility that implicit adaptation only occurred after participants had been aware of the temporal regularity in the preceding explicit phase. In sum, these results indicate that participants not only adapted to the regular target intentionally (explicit phase), but also implicitly without detecting its presence (implicit phase).

## Experiment 2

The results from Experiment 1 revealed implicit adaptation to the ‘hidden’ temporal regularity. However, in-depth analyses of the pseudorandomly generated stimulus sequences revealed that the average inter-stimulus interval differed between regular and irregular targets. To ensure that none of the observed effects could be attributed to this inadvertent higher-order dependency, we replicated Experiment 1 with a more stringent procedure to generate stimulus sequences.^3^

### Method

Power analyses based on RT effects in Experiment 1 indicated that data from 32 participants would suffice to again detect an effect of ‘regularity’ (Cohen’s *d* = 0.51, power = 80 %, *α* = 0.05). Yet, as Experiment 2 served as a stringent replication of Experiment 1 we collected data until 48 participants were included using the same ‘Inclusion Criteria’ as for Experiment 1, maintaining full counterbalancing of experimental conditions. In total, 54 participants participated for course credits (six excluded). The experimental procedure was approved by the Ethical committee of the Faculty of Behavioural and Social Sciences (18233-S), and participants gave written informed consent prior to testing.

### Results

#### Questionnaire

None of the participants indicated awareness of the regularity when asked whether they had noticed anything ‘remarkable’. In the yes/no question, five out of 54 participants (9.3 %) indicated to have been aware of the regularity. In the identification task, 23 participants (42.6 %) indicated the correct location of the regular target. Of those, two also indicated being aware of the regularity and therefore were excluded from further analyses.

#### Reaction times

Figure 3a and b show similar RT results to Experiment 1. Indeed, statistical analyses supported the inclusion of the factors ‘regularity’ *(ΔBIC* = 281.54, *BF* > 1000; *χ*^2^(3) = 312.99, *p* < 0.001), ‘phase’ (Δ*BIC* = 242.11, *BF* > 1000; *χ*^2^(2) = 263.07, *p* < 0.001), and their interaction *(ΔBIC* = 221.72, *BF* > 1000; *χ*^2^(1) = 232.21, *p* < 0.001). The interaction entailed that regular and irregular targets differed more in the explicit than the implicit phase. Post hoc Tukey’s HSD test revealed that RTs to the regular target were significantly lower in both the explicit (*z* = 18.70, *p* < 0.001) and implicit phase (*z* = 5.26, *p* < 0.001). Additionally, we again found no evidence for an effect of the time course on adaptation *(ΔBIC* = 19.99, *1/BF* > 1000; *χ*^2^(4) = 6.46, *p* = 0.17).

**Figure 3.**
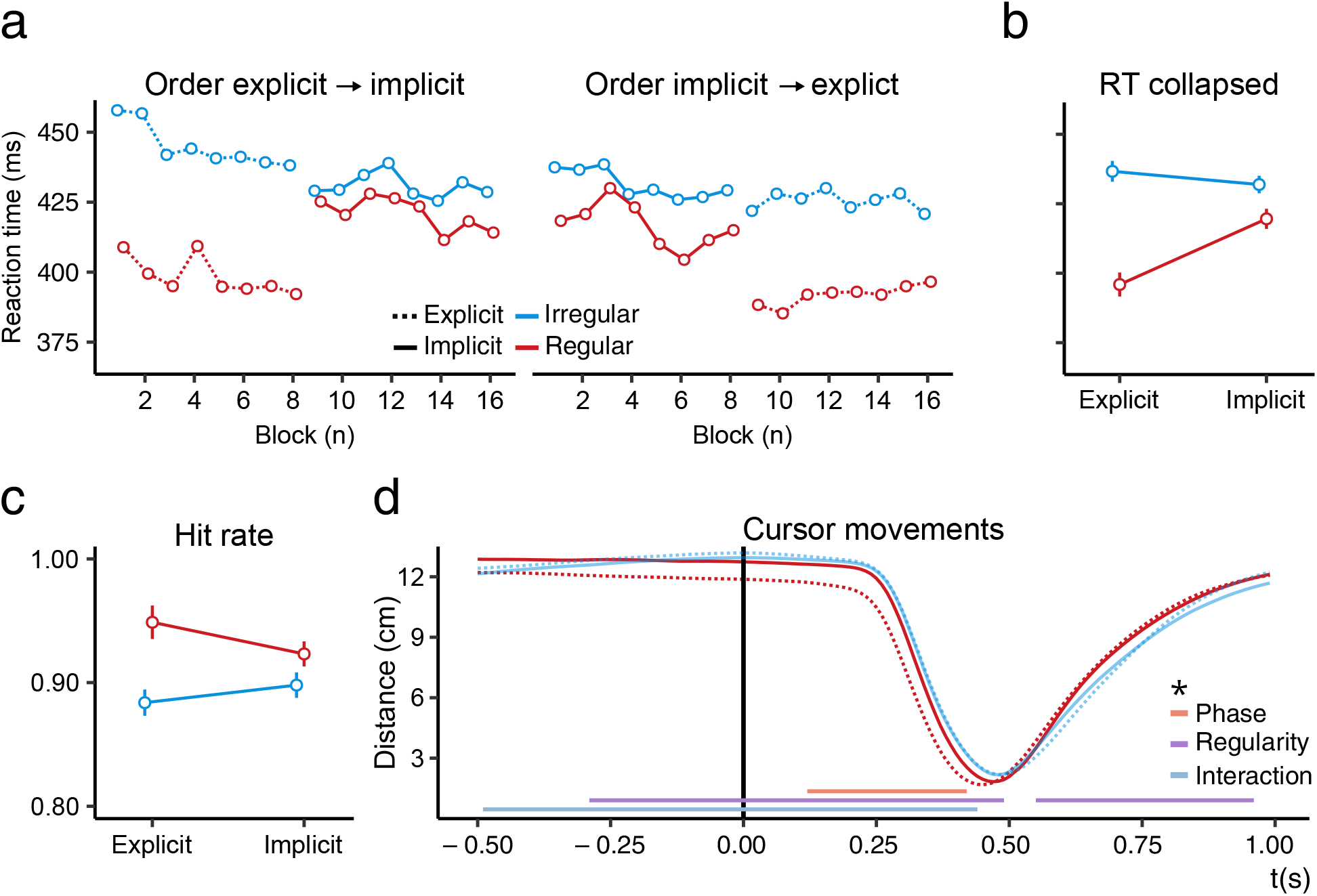
Results of Experiment 2. Reaction time (a and b), hit rate (c), and cursor movements (d). Plotted as in Figure 2.

#### Hit rate

Results regarding hit rate (Figure 3c) were similar to Experiment 1. GLMM comparisons revealed support for the inclusion of ‘regularity’ *(ΔBIC* = 58.62, *BF* > 1000; *χ*^2^(3) = 90.35, *p* < 0.001) and no support for an effect of time course on adaptation (Δ*BIC* = 23.05, 1/*BF* > 1000; *χ*^2^(4) = 3.40, *p* = 0.49). Unlike Experiment 1, here we found support for the inclusion of ‘phase’ interacting with ‘regularity’ *(ΔBIC* = 31.26, *BF* > 1000; *χ*^2^(2) = 52.41, *p* < 0.001). The interaction shows that the difference between the regular and irregular targets was more pronounced in the explicit phase. Post hoc Tukey’s HSD test revealed that hit rate was nevertheless significantly higher to the regular target in both the explicit *(z* = −10.03, *p* < 0.001) and implicit phase *(z* = −3.48, *p* < 0.001).

#### Cursor movements

In Experiment 2, cursor movements revealed significant effects of ‘regularity’, ‘phase’, and their interaction (Figure 3d). Two significant clusters for ‘regularity’ were found from 0.29 s before to 0.49 s after target onset (*p* < 0.001) and 0.55 s to 0.96 s after target onset (*p* < 0.001), similar to Experiment 1. A main effect of ‘phase’ was marked by a significant cluster from 0.12 s to 0.42 s after target onset (*p* = 0.02), and a cluster for their interaction was found from 0.49 s before to 0.44 s after target onset (*p* < 0.001). This interaction entailed a stronger bias towards regular targets in the explicit phase compared to the implicit phase.

#### Conflict situations

We pre-registered an additional hypothesis regarding hit rates (https://tinyurl.com/w3ocv8x) in ‘conflict situations’, where two targets are simultaneously on screen. We expected that in such situations participants would be more inclined to hit the regular target. We found no support for this hypothesis, neither in the explicit or implicit phase (see Supplemental Material for the statistical report). Conflict situations were rare (~5 % of trials), so this was possibly due to a lack of power.

### Discussion

Experiment 2 served as a stringent replication of Experiment 1, and ascertained that the adaptation effects were exclusively caused by the temporal regularity. All crucial effects of Experiment 1 were replicated: RT was found to be lower, hit rate to be higher, and the cursor movements to be initiated earlier to the regular targets compared to irregular targets. Furthermore, we found an interaction between ‘regularity’ and ‘phase’ in the hit rate and cursor movement analyses, indicating a stronger bias towards regular targets in the explicit than implicit phase (Figure 3c and d).

## General discussion

Do we make use of regular intervals in dynamic, complex environments to decide when to act even when we are oblivious to them? To answer this question, we developed a game-like experimental paradigm where participants scored points by ‘hitting’ sudden-onset targets presented in random order. One of these targets appeared every three seconds, the others with random timing. Across two experiments, less than 5 % of the participants detected this regularity. Nevertheless, we found that unaware participants still adapted to the regularity, as indicated by RT, hit rates, and cursor movements. The high hit rate to both regular and irregular targets show that this implicit adaptation occurred even though responses to the regularity were interrupted by responses to irregular events. In sum, participants adapted to the temporal regularity without detecting its presence, and thus without intentionally utilizing temporal information.

Previous research on temporal preparation already suggested that participants adapt to temporal regularities. In these studies, timing is often characterized as ‘implicit’ since participants are not required to make an overt time estimation (see Coull & Nobre, 2008, for a taxonomy). However, each trial presents an isolated and explicitly cued interval, and therefore the relevance of temporal information for the task is transparent. This contrasts with everyday behavior in which we learn to act based on complex, uncued, dynamically unfolding events and are hardly ever aware of the temporality of our behavior. A critical question is whether findings from timing research generalize to naturalistic environments (van Rijn, 2018). The challenge in such less controlled environments is to maintain the precision and rigorous methods of laboratory experiments. Our game-like ‘Whac-A-Mole’ experiment is an example of how laboratory environments can be designed to measure temporal behavior under naturalistic conditions (see also Kunchulia, Parkosadze, & Thomaschke, 2019; Schlichting et al., 2018; van Rijn, 2014).

A recent study from our group (Damsma, Taatgen, de Jong, & van Rijn, 2019) also studied adaptation to implicit temporal regularities. In their experiments, a visual search display was preceded by sequences of stimuli, and one location or feature in this sequence was presented at regular intervals. They hypothesized that attention would be attracted by the temporal regularity, but no evidence for such adaptation was found. A critical difference with the present study is that in our paradigm the regularity was part of the task’s goal, thereby immediately pairing the regular target with a motor response and with rewarding feedback upon a successful hit. Previous research has demonstrated that motor learning is facilitated when predictive cues are directly associated with a motor plan and immediate reward (Howard, Wolpert, & Franklin, 2015; Manohar et al., 2015; Wolpert & Flanagan, 2016). It is likely that in our design participants therefore attended the regularity more than in Damsma et al. (2019), which has previously been argued to be crucial for statistical learning (Baker et al., 2004; Richter & de Lange, 2019; Turk-Browne et al., 2005).

Research on motor learning of rhythmic sequences has similarly stressed the tight link between timing and motor action in implicit learning (Gobel et al., 2011; O’Reilly et al., 2008; Shin & Ivry, 2002). Typically, in these studies, often using a version of the serial reaction time task, a fixed sequence of finger movements is matched with a rhythmic stimulus sequence (e.g., stimuli ‘A’, ‘B’, and ‘C’ are responded to with finger ‘1’, ‘2’, and ‘3’). Participants implicitly adapt to the intervals between each subsequent movement in the motor sequence, thereby decreasing their RT. However, when decoupling the temporal structure from the response sequence by randomizing stimulus order, such adaptation did not occur. These findings suggested that temporal information was not learned independently from the ordinal sequence of motor actions. Our findings challenge this conclusion. The regularity in our study was decoupled from the motor sequence order by embedding it in an otherwise irregular stream of events, yet participants implicitly adapted to the regularity. This discrepancy might be ascribed to the relative simplicity of our three second regular interval compared to regularities consisting of different durations used in these motor learning studies. Alternatively, although our regularity was decoupled from the ordinal motor sequence, the location of the regularity was fixed and therefore coupled to the movement direction. Thus, while a prerequisite for learning of timing information might be that the regularity is to some extent coupled to motor action, it may not necessarily have to be coupled with an ordinal sequence of motor actions.

What mechanism could give rise to such implicit adaptation to temporal regularities? A potential neural correlate is rhythmic neural entrainment, a process where neural oscillations align to the temporal structure of a stimulus sequence (Large & Jones, 1999; Large & Palmer, 2002; Schroeder & Lakatos, 2009). Interestingly, Herbst and Obleser (2017, 2019) recently investigated such entrainment in a timing task in which a warning cue predicted the temporal onset of the target. Results indicated that oscillations in the delta band (1-3 Hz) implement temporal predictions by a phase-locking mechanism that aligns more excitable phases of neural oscillations with the target’s onset. This phase-locking mechanism was initiated by a warning cue. An outstanding question is whether such entrainment can also arise in the absence of explicit warning cues to give rise to implicit learning of temporal behavior in a natural setting. Our paradigm would enable one to study the role of neural entrainment in implicit learning of temporal behavior in a natural, dynamic setting.

Such a phase-locking mechanism could account for our results if the temporal predictions formed by such entrainment would, for example, lead to enhancement of temporal attention around the moment of regular target onset. It should be noted, however, that our design does not allow one to separate enhancement of responses to the regular target versus active suppression of responses to the irregular targets. Although participants were instructed and encouraged to divide their attention equally between all targets, it may be inevitable that implicit prioritization of the regular target comes at the cost of performance to the irregular targets. Yu and Zhao (2015) investigated this particular dissociation for implicit adaptation to regularities in stimulus order, rather than in their timing. Their results suggested that adaptation was expressed as attentional enhancement of the regularity rather than a suppression of irregular targets. It could be interesting to explore whether their approach could be adopted to determine the exact nature of adaptation to regular intervals too.

### Conclusion

We aimed to measure naturalistic timing behavior through a paradigm in which a regular interval is embedded in a stream of irregularly timed events. We found that participants automatically adapted to this regularity, without being able to report it afterwards. In this way, we provide direct evidence that regularities in the time between the events we encounter in our daily life can be used to decide when to act, whilst remaining undetected.

## Open Practices Statement

The data and materials for all experiments are available at (https://tinyurl.com/w3ocv8x) and Experiment 2 was preregistered (https://osf.io/hdycr/).

## Appendix

### Questionnaire

Figure A1 displays the self-report measure to assess participant’s awareness of the temporal regularity. As displayed, in question 1 we first encouraged participants to report anything they might have found ‘remarkable’ in the experiment, thereby stimulating false-positive responses. Only ~4 % of the participants reported the presence of the regularity in this open question. In question 2, we explicitly informed participants about the regularity, and asked them whether they had noticed this. In question 3, participants were asked to indicate the location of the regular target. In this way, we assessed whether participants that indicated in question 2 to be aware of the regularity could also correctly locate the regularity.

**Figure A1.**
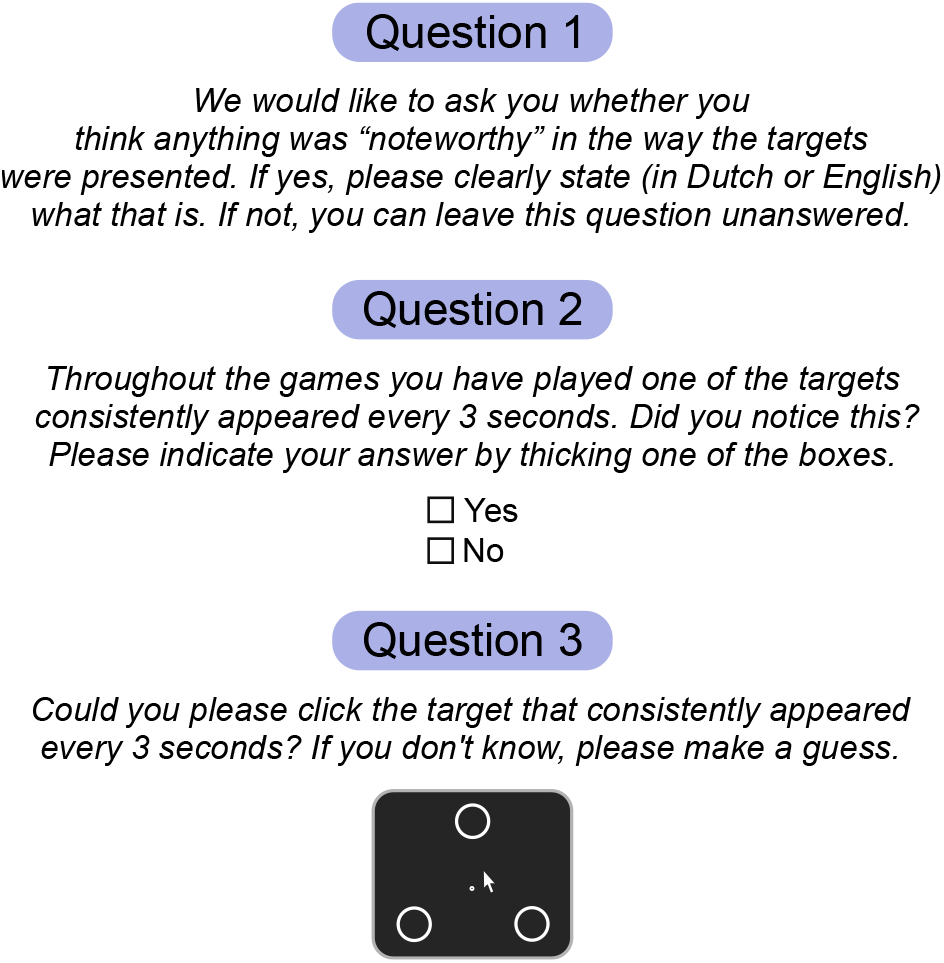
The complete questions used to assess participants’ awareness of the regularity after the implicit phase, as presented to the participants with phase order: ‘implicit → explicit’. The group ‘explicit → implicit’ saw the same questions except that question 2 was phrased: *‘Although, we told you that the temporal regularity would be removed in the second part of the experiment, there was still one target consistently presented every three seconds. Did you notice this?’*

### Stimulus sequences

Experiment 2 was a direct, more stringent replication of Experiment 1, and was conducted because we found that the inter-stimulus interval (ISI) in Experiment 1 differed for regular versus irregular targets. To rule out that the observed effects might have been driven by this higher-order dependency, we replicated Experiment 1 with a more stringent randomization procedure for the stimulus sequences. In order to minimize ISI differences in conditions while otherwise maintaining the randomization procedure as unconstrained as in Experiment 1, we generated stimulus sequences for each participant for each block in the experiment and assessed the ISI distributions for regular and irregular targets by means of the Kullback-Leibler (KL) divergence. That is, for each stimulus sequence we estimated the ISI distributions for both conditions using a Gaussian kernel density estimation with a fixed bandwidth of 0.2, and computed the KL-divergence as a measure of how one probability distribution (regular) diverges from a second probability distribution (irregular). A sequence was only accepted as an experimental sequence if the KL-divergence between the two ISI distributions was lower than 0.001 (for comparison, in Experiment 1 the KL-divergence was ~0.1). Table A1 provides a descriptive summary of the ISI distributions in both experiments. The code used to generate these sequences is included in the OSF-repository (https://tinyurl.com/w3ocv8x).

**Table A1.**
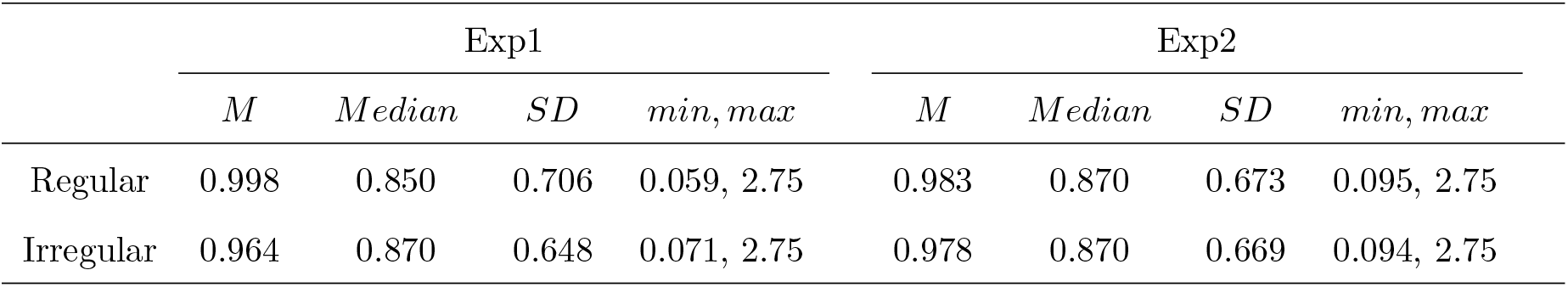
Descriptive summary of the ISI distributions of Experiment 1 (Exp1) and Experiment 2 (Exp2): mean (M), median, standard deviation (SD), and minimum and maximum ISI (min, max), all displayed in seconds.

|  | Exp1 |  |  |  | Exp2 |  |  |  |
| --- | --- | --- | --- | --- | --- | --- | --- | --- |
|  | *M* | *Median* | *SD* | *min, max* | *M* | *Median* | *SD* | *min, max* |
| Regular | 0.998 | 0.850 | 0.706 | 0.059, 2.75 | 0.983 | 0.870 | 0.673 | 0.095, 2.75 |
| Irregular | 0.964 | 0.870 | 0.648 | 0.071, 2.75 | 0.978 | 0.870 | 0.669 | 0.094, 2.75 |

## Footnotes

1 Table A1 in the Appendix provides a full report on the descriptive statistics of the generated intervals: the mean, median, standard deviation, minimum, and maximum.

2 Analyses without excluding direct repetitions yielded the same conclusions.

3 In the Appendix, we discuss this issue in more detail as well as the measures taken against it. For Experiment 2, the hypotheses and analysis plan were pre-registered: https://tinyurl.com/w3ocv8x. In addition to these pre-registered analyses, we include analyses of the effect of time course on adaptation (see Figure 2a and 3a) which were suggested by an anonymous reviewer.

