## Supplemental Materials for "Implicit Learning of Temporal Behavior in Complex Dynamic Environments"

### Supplemental Material: Implicit Learning of Temporal Behavior in Complex Dynamic Environments

Author Note

Correspondence concerning this Supplemental document should be addressed to Josh M. Salet, Department of Experimental Psychology, University of Groningen, Grote Kruisstraat 2/1, 9712 TS Groningen, the Netherlands.

Supplemental Material: Implicit Learning of Temporal Behavior in Complex Dynamic Environments

#### S.1. Statistical report

In this section, we elaborate on the statistical procedure for the model selection of the (General) Linear Mixed Models (G)LMM to analyze response time (RT) and hit rate. In addition, we report the full fixed- and random effects structure of the models. In order to report classical effect sizes, we ran a repeated measures analyses of variance (ANOVA). Its results similarly support the conclusions of our (G)LMM analyses.

Statistical evidence for or against an effect of interest was quantified by comparing the best statistical model (in terms of $BIC$) with this effect, against the best model without this predictor term. For some comparisons, this means that the compared models can differ in terms of other predictors as well. For example, when testing for an interaction effect $A \times B$ on RT, the best model *with* might be $RT \sim A \times B$, whereas the best model without this interaction might be $RT \sim A$, which omits the main effect of $B$ as well. In the R-notebooks on our OSF repository (https://tinyurl.com/w3ocv8x) we list full tables with all possible models ranked on their BIC, which determines which models were actually compared.

##### Response time and hit rate

**Response time.** In addition to the support for the fixed effects reported in our article ('regularity', 'phase', and their interaction), there was support for the fixed effect 'previous response type' (hit/miss) in both Experiment 1 ($\Delta BIC = 5.85$, $BF = 18.64$; $\chi^2(1) = 16.14$, $p < 0.001$) and Experiment 2 ($\Delta BIC = 13.44$, $BF = 827.29$; $\chi^2(1) = 23.92$, $p < 0.001$). RTs were higher when the preceding trial was a miss, compared to when it was a hit, $M$: 0.42 s and 0.41 s (Experiment 1); 0.44 s and 0.41 s (Experiment 2). In both Experiments, the random effects structure of the model included, besides an intercept for each participant, a random slope for the effect of 'regularity' and 'previous response type'.

Table S1

*ANOVA table for RTs in Experiment 1 (Exp1) and Experiment 2 (Exp2).*

| | Exp1 | | | | Exp2 | | | |
|---|---|---|---|---|---|---|---|---|
| | $df1, df2$ | $F$ | $p$ | $\eta_p^2$ | $df1, df2$ | $F$ | $p$ | $\eta_p^2$ |
| regularity | 1, 39 | 59.50 | < 0.001 | 0.60 | 1, 47 | 176.69 | < 0.001 | 0.79 |
| phase | 1, 39 | 10.48 | 0.002 | 0.21 | 1, 47 | 8.46 | 0.006 | 0.15 |
| regularity * phase | 1, 39 | 10.14 | 0.003 | 0.21 | 1, 47 | 32.82 | < 0.001 | 0.41 |

We performed a two-way repeated-measures ANOVA regarding our main interests: 'regularity', 'phase', and their interaction. This yielded the same conclusions for both Experiments (see Table S1). RT was lower for the regular target compared to the irregular targets, lower in the explicit phase than implicit phase, and the interaction between 'regularity' and 'phase' revealed that regular and irregular targets differed more in the explicit than the implicit phase.

**Hit rate.** The GLMM analyses on hit rate revealed that, besides the support for the fixed effects reported in the main text (Experiment 1: 'regularity'; Experiment 2: 'regularity', 'phase', and their interaction), there was support for the inclusion of a fixed effect of 'previous response type' in both Experiment 1 ($\Delta BIC$ = 80.36, $BF >$ 1000; $\chi^2(1)$ = 90.73, $p <$ 0.001) and Experiment 2 ($\Delta BIC$ = 38.91, $BF >$ 1000; $\chi^2(1)$ = 49.49, $p <$ 0.001). The probability of a hit was higher in case of a hit on the previous trial compared to a miss on the previous trial, $M$: 0.94 and 0.90 (Experiment 1); 0.94 and 0.88 (Experiment 2). In both Experiments, the random effects structure of the models included a random intercept for each participant and a random slope term for the effect of 'regularity'. In Experiment 2, the random effects structure additionally included terms for 'phase', and 'previous response type'

A two-way repeated-measures ANOVA with the factors 'regularity' and 'phase' and their interaction revealed a main effect of 'regularity', and the interaction between 'regularity' and 'phase', but not for the main effect of 'phase' (see Table S2).

Table S2

*Same as in Table S1, but for hit rates.*

| | Exp1 | | | | Exp2 | | | |
|---|---|---|---|---|---|---|---|---|
| | $df1, df2$ | $F$ | $p$ | $\eta_p^2$ | $df1, df2$ | $F$ | $p$ | $\eta_p^2$ |
| regularity | 1, 39 | 11.42 | 0.002 | 0.23 | 1, 47 | 42.79 | $< 0.001$ | 0.48 |
| phase | 1, 39 | 0.06 | 0.80 | 0.002 | 1, 47 | 0.27 | 0.60 | 0.01 |
| regularity * phase | 1, 39 | 5.57 | 0.02 | 0.13 | 1, 47 | 19.01 | $< 0.001$ | 0.29 |

#### Phase order

As indicated in the main text, model comparison in Experiment 1 and 2 did not support the inclusion of the main effect 'phase order' for either RT (Table S3) or hit rate (Table S4). To further investigate order effects, we assessed whether 'phase order' interacted with the main predictors 'regularity' and 'phase'. These analyses did not reveal any effect on either RT or hit rate (Table S3 and S4). Note that Bayesian Inference did not always correspond to the outcome of Likelihood Ratio Tests, reflecting the conservative nature, of Bayesian tests penalizing model complexity more severely. Although we report the outcome of the Likelihood Ratio Tests for completeness, we relied on the *BIC*, as a prespecified criterion, for our conclusions.

**Time course on adaptation.** To analyze whether the time course within each phase affected adaptation (expressed by RT and hit rate) we constructed a separate LMM.

Table S3

*Bayesian inference ($\Delta BIC$, $1/BF$) and Likelihood Ratio Tests ($\chi^2$, $df$, and $p$) for RTs, assessing the effect of 'phase order' on 'regularity', 'phase', and their interaction.*

| | Exp1 | | | | | Exp2 | | | | |
|---|---|---|---|---|---|---|---|---|---|---|
| | $\Delta BIC$ | $1/BF$ | $\chi^2$ | $df$ | $p$ | $\Delta BIC$ | $1/BF$ | $\chi^2$ | $df$ | $p$ |
| phase order | 9.75 | 130.86 | 0.54 | 1 | 0.46 | 10.40 | 181.59 | 0.08 | 1 | 0.78 |
| regularity * phase order | 19.38 | $> 1000$ | 1.19 | 2 | 0.55 | 20.72 | $> 1000$ | 0.25 | 2 | 0.88 |
| phase * phase order | 5.77 | 17.88 | 14.80 | 2 | $< 0.001$ | 9.91 | 141.88 | 11.06 | 2 | 0.004 |
| regularity * phase * order | 13.33 | 782.93 | 27.81 | 4 | $< 0.001$ | 12.93 | 643.25 | 28.99 | 4 | $< 0.001$ |

Table S4

*Same as in Table S3, but for hit rates.*

| | Exp1 | | | | | Exp2 | | | | |
|---|---|---|---|---|---|---|---|---|---|---|
| | $\Delta BIC$ | $1/BF$ | $\chi^2$ | *df* | *p* | $\Delta BIC$ | $1/BF$ | $\chi^2$ | *df* | *p* |
| phase order | 10.32 | 174.44 | 0.05 | 1 | 0.83 | 9.56 | 118.92 | 1.02 | 1 | 0.31 |
| regularity * phase order | 19.84 | $> 1000$ | 0.90 | 2 | 0.64 | 17.86 | $> 1000$ | 3.29 | 2 | 0.19 |
| phase * phase order | 20.51 | $> 1000$ | 10.60 | 3 | 0.01 | 20.07 | $> 1000$ | 1.08 | 2 | 0.58 |
| regularity * phase * order | 30.66 | $> 1000$ | 31.56 | 6 | $< 0.001$ | 24.18 | $> 1000$ | 18.12 | 4 | 0.001 |

We first determined the average RT and hit rate difference between regular and irregular targets for each experimental block (‘block’) split on ‘phase’ and ‘phase order’. Subsequently, we re-coded the predictor ‘phase’ and ‘phase order’ in the single predictor ‘all order’ with four levels: ‘Explicit - Phase 1’, ‘Explicit - Phase 2’, ‘Implicit - Phase 1’, and ‘Implicit - Phase 2’. Each of these four phases comprised eight experimental blocks. We then compared LMMs to test for the effect of ‘block’ (as continuous predictor), ‘all order’, and their interaction. The random effects structure in both experiments, for both RT and hit rate, included a random intercept for each participant and a random slope term for the effect of ‘all order’.

Table S5

*Bayesian inference ($\Delta BIC$, $1/BF$) and Likelihood Ratio Tests ($\chi^2$, $df$, and $p$) for the effect of time course on adaptation expressed by the difference in RT for regular and irregular targets, assessing the effect of ‘block’, ‘all order’, and their interaction. Note that all Bayes factors indicate the absence of an effect ($1/BF$), except for the effect of ’all order’ in row 2 ($BF$)*

| | Exp1 | | | | | Exp2 | | | | |
|---|---|---|---|---|---|---|---|---|---|---|
| | $\Delta BIC$ | $1/BF$ | $\chi^2$ | *df* | *p* | $\Delta BIC$ | $1/BF$ | $\chi^2$ | *df* | *p* |
| block | 5.57 | 16.23 | 0.83 | 1 | 0.36 | 6.27 | 23.02 | 0.34 | 1 | 0.56 |
| all order | 8.79 | 81.05 ($= BF$) | 10.43 | 3 | 0.015 | 6.35 | 23.89 ($= BF$) | 26.18 | 3 | $< 0.001$ |
| block * all order | 19.75 | $> 1000$ | 5.88 | 4 | 0.21 | 19.99 | $> 1000$ | 6.46 | 4 | 0.17 |

Table S6

*Same as in Table S5, but for hit rates.*

| | Exp1 | | | | | Exp2 | | | | |
|---|---|---|---|---|---|---|---|---|---|---|
| | $\Delta BIC$ | $1/BF$ | $\chi^2$ | $df$ | $p$ | $\Delta BIC$ | $1/BF$ | $\chi^2$ | $df$ | $p$ |
| block | 5.08 | 12.72 | 1.32 | 1 | 0.25 | 4.59 | 9.91 | 2.02 | 1 | 0.16 |
| all order | 4.63 | 10.15 | 14.59 | 3 | 0.002 | 31.07 | $> 1000$ ($= BF$) | 50.90 | 3 | $< 0.001$ |
| block * all order | 23.75 | $> 1000$ | 21.10 | 7 | 0.003 | 23.05 | $> 1000$ | 3.40 | 4 | 0.49 |

As displayed in Table S5, model comparisons did not show support for an effect of 'block', neither as main effect nor as interacting term with 'all order'. However, in both experiments, model comparisons supported the inclusion of 'all order'. Post hoc Tukey's HSD tests revealed that the difference in RT between regular and irregular targets in Experiment 2 was more expressed in the explicit than implicit phase, regardless of the order. Instead, in Experiment 1, this was only true for 'Explicit - Phase 2' compared to 'Implicit - Phase 1' ($t$ = -2.84, $p$ = 0.035).

The statistical report for the average hit rate difference is displayed in Table S6. Model comparisons did not show support for an effect of 'block' in both experiments. In Experiment 2, model comparisons supported the inclusion of 'all order', but not in Experiment 1. Post hoc Tukey's HSD tests revealed that the difference in hit rate between regular and irregular targets in Experiment 2 was larger for the level 'Explicit - Phase 1' compared to levels 'Implicit - Phase 1' ($t$ = 3.78 , $p$ = 0.002) and 'Implicit - Phase 2' ($t$ = 6.77 , $p < 0.001$).

**Conflict Situations.** For Experiment 2 we pre-registered an additional hypothesis regarding hit rates in conflict situations, that is, situations with near-simultaneously presented targets (https://tinyurl.com/w3ocv8x). We hypothesized that in such situations participants might be more likely to miss (or skip) irregular targets if a predictable regular target was to be presented while the irregular target was still on screen. To test this, we selected only irregular targets $T_n$ with a subsequent $T_{n+1}$ appearing within 0.5 s. Note that each target was on screen for 0.5 s. We then contrasted hit rates on $T_n$ separately for $T_{n+1}$

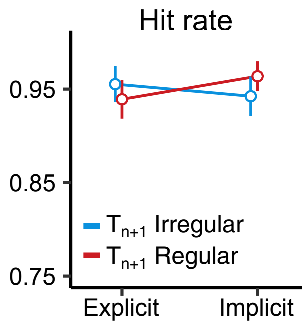


*Figure S1*. Hit rate in conflict situations. The data points display the hit rate to $T_n$ split on $T_{n+1}$ = regular and $T_{n+1}$ = irregular. The plot is based on a subset of the data excluding three participants that did not have data for all conditions.

= regular and $T_{n+1}$ = irregular. In this analysis, we only used data where $T_{n+1}$ was hit, and Tn appeared more than 0.25 s after its preceding $T_{n-1}$ (to prevent situations of triple conflict). This selection procedure resulted in a very small subset of the data (∼5 %).

Figure S1 displays the hit rate in conflict situations. If participants would prioritize a regular $T_{n+1}$, it is expected that the hit rate to $T_n$ is lower when $T_{n+1}$ is regular, compared to irregular. Such a trend is not present. Indeed, (G)LMM analyses did not reveal support for any of the tested predictors of interest: '$T_{n+1}$ regularity' ($\Delta BIC = 7.59$, $1/BF = 44.41$; $\chi^2(1) = 0.02$, $p = 0.90$), 'phase' ($\Delta BIC = 5.58$, $1/BF = 16.25$; $\chi^2(1) = 2.02$, $p = 0.15$), and their interaction ($\Delta BIC = 18.07$, $1/BF > 1000$; $\chi^2(3) = 4.74$, $p = 0.19$).

Using a two-way repeated-measures ANOVA with the factors '$T_{n+1}$ regularity' and 'phase' we arrived at the same conclusions. None of the predictors reached significance: '$T_{n+1}$ regularity' ($F(1, 44) = 0.04$, $p = 0.85$, $\eta_p^2 < 0.001$), 'phase' ($F(1, 44) = 0.19$, $p = 0.66$, $\eta_p^2 = 0.004$), and their interaction ($F(1, 44) = 1.78$, $p = 0.19$, $\eta_p^2 = 0.04$). Note that after the selection procedure to obtain the conflict situations, three participants did not have data for all levels (split on 'phase' and 'regularity'). The ANOVA was performed on the subset of the data excluding these three participants.
